# Theta phase precession at encoding predicts subsequent memory of sensory-driven vector fields, & occurs in memory-dependent fields at retrieval

**DOI:** 10.1101/2023.06.05.543704

**Authors:** Steven Poulter, William de Cothi, Caswell Barry, Colin Lever

## Abstract

Theta phase precession is thought to confer key computational advantages (e.g. temporal compression suiting spike-timing related plasticity, cognitive relations as phase distances, and population-level coding for directions and sequences). However, direct evidence speaking to: 1) its widely-theorised role in enhancing memorability; 2) its dependence upon sensory input, is lacking. We leveraged the Vector trace cell (VTC) phenomenon to examine these issues. VTCs in subiculum show a simple, unambiguous memory correlate: VTCs remember the distance and direction to a cue after the cue is removed, with a new ‘trace field’ which was not present before the cue was inserted. Regarding memorability, here we show that subsequently-remembered cue fields (those which become trace fields) exhibit higher levels of phase precession than subsequently-forgotten cue fields (those which produce no trace). Thus, phase precession does appear to enhance memorability, consistent with long-established theory. The second issue concerns the extent of phase precession in sensory-elicited vs memory-dependent firing. Phase precession in CA1 is strongly disrupted following deprivation of its Entorhinal, but not CA3, inputs; this could indicate that theta phase precession is largely sensory-driven and absent in memory-dependent fields. Here, however, we show that phase precession is robust in subicular VTC trace fields, i.e. when the cue that originally elicited the new vector field is no longer present. Thus, the much-theorised benefits of phase precession likely apply to memory-dependent fields. These findings have wide implications for oscillatory-based models of memory.

## Introduction

The hippocampal formation is widely thought to contribute to representations of space (O’Keefe, 1976; O’Keefe and Nadel, 1978; Poulter et al, 2018) and of time (Pastalkova et al, 2008; Kraus et al, 2013; Eichenbaum, 2014). For example, one hippocampal CA1 cell might fire when a rodent is at a specific location in a spatial context or at a specific temporal stage in a task, while another CA1 cell fires at a different location or temporal stage. A robust and theoretically-provocative phenomenon called ‘theta phase precession’ occurs during traversal of such fields, linking these cognitive variables (place and time) to the large amplitude, 6-12Hz theta oscillation (Buzsaki, 2006) in the hippocampus. Phase precession refers to the common observation that as the animal progresses through the spatial or temporal field, the CA1 cell fires at successively earlier stages of each theta cycle (O’Keefe and Recce, 1993; Pastalkova et al, 2008; Shimbo et al, 2021) over the ∼4-9 theta cycles of the firing field.

Phase precession not only tightly links cognitive to oscillatory variables, it is also a remarkably pervasive phenomenon, and is thus particularly important to understand. 1) As summarised above, it occurs for both spatial and temporal variables. 2) While theta phase precession was first noticed in hippocampal place cells on linear tracks (O’Keefe and Recce, 1993; Skaggs et al, 1996), it also occurs in hippocampal place cells and entorhinal grid cells in the open field (Huxter et al, 2008; Climer et al, 2013; Jeewajee et al, 2014). 3) Theta phase precession occurs not only in different regions *within* the hippocampal formation (O’Keefe and Recce, 1993; Hafting et al, 2008; Kim et al, 2012, Lee et al, 2022), but also in various regions *beyond* it (e.g. Jones & Wilson, 2005; Van Der Meer and Redish, 2011), suggesting its brain-wide application and relevance to several adaptive functions. 4) During high-frequency (140-200 Hz) ripples, CA1 place cell spiking has recently been to shown to precess relative to the ripple oscillation (Bush et al, 2022). Accordingly, phase precession may reflect general neural coding principles transcending any one oscillatory band (Bush & Burgess, 2020). 5) Theta and other-band phase precession has been found in the human brain (Qasim et al, 2021), suggesting that revealing phase precession mechanisms and functional associations in the rodent will enlighten human neurocognitive processes.

Thus, theta phase precession is remarkably pervasive, and merits empirical examination of the functions that it confers.

### The computational advantages of theta phase precession: enhancing memorability?

Theta phase precession has been theorised to represent relationships between cognitive variables (e.g. field B’s peak is 40cm east of field A’s peak) as phase distances (cell B fires 50 degrees later than A in the theta cycle in eastwards motion), with theta-compressed representations suiting spike-timing related plasticity (e.g. Burgess et al, 1994; Skaggs et al, 1996; Lisman, 2005; Dragoi & Buzsaki, 2006). A common view is that theta phase precession enables population-level coding for sequences (e.g. Skaggs et al, 1996; Lisman, 2005; Dragoi & Buzsaki, 2006) and directions (e.g. Huxter et al, 2008; Zutshi et al, 2017; Bush & Burgess, 2020).

A shared theme running through these and other theoretical models of theta phase precession is that phase precession aids learning. For instance, in Burgess et al, 1994, a navigation network exploits phase coding to remember the direction to a goal. Perhaps the most dominant view of theta phase precession is that it supports sequence learning. In models of sequence learning (e.g. Skaggs et al, 1996; Lisman, 2005; Dragoi & Buzsaki, 2006), ordinal relationships experienced on a seconds-timescale are preserved, but also compressed, by theta phase precession into a millisecond timescale suitable for well-established plasticity regimes such as spike-timing dependent long term plasticity (STDP: Bi and Poo, 1998). The basic concept is that cells with similarly-sized fields showing theta phase precession will tend to fire in a theta phase-dependent orderly sequence. Within a given theta cycle, a set of A cells firing at earlier phases tend to fire before B cells firing at middle phases, which in turn tend to fire before C cells firing at later phases. These phase offsets translate into pre-before-postsynaptic temporal intervals (e.g. B cells fire ∼25ms after A cells, C cells fire ∼25ms after B cells) that are well-suited, over repeated exposure, for asymmetric STDP to ‘stamp in’ an A->B->C spatial or temporal sequence. After synaptic plasticity occurs, even in the absence of the sensory stimuli associated with B cells, A cells will start to make B cells fire, and thus predict B type associations. An analogy is that on first exposure(s) to an album/playlist, one does not hear the beginning of song 2 in ‘the mind’s ear’ after song 1 ends, but this predictive ability does occur after a few plays of that auditory experience. Thus, theta phase precession may enable sequence learning and predictive coding (e.g. Mehta et al, 1997; Lisman & Redish, 2009; Stachenfeld et al, 2017; Reifenstein et al, 2021; George et al, 2023).

Explicit computational modelling of the learning of sequential order suggests that important contributions are played by *both* phase precession *and* asymmetric spike-timing plasticity rules (e.g. Reifenstein et al, 2021). However, the relative importance of symmetric vs asymmetric timing induction rules, and the general applicability of STDP in behaving animals remains unclear, and plasticity regimes other than STDP may also turn out to be important in real-world memory (Bittner et al, 2017). Accordingly, then, for several reasons it is crucial to empirically test if theta phase precession predicts higher levels of memorability.

Up till now, it has been difficult to conduct such tests at least in part because unambiguously mnemonic correlates of hippocampal cell firing have arguably been elusive at the single-cell level. In our view, subicular Vector Trace cells (VTCs) provide a useful model. The insertion of a cue object such as a barrier or wine bottle elicits a new firing field in a VTC and that firing field persists, after the cue object is removed, in the form of a trace field (Fig 1a). This provides an opportunity to explore to what extent the presence of phase precession (Fig 1b) in the cue field (encoding trial) predicts the likelihood of a trace field (subsequent memory), and thus to address our first question, ‘Does phase precession enhance memorability?’ (Fig 1c). It is possible that some of form of phase-precession-supported sequence learning is part of the mechanism for generating trace fields.

**Fig. 1.**
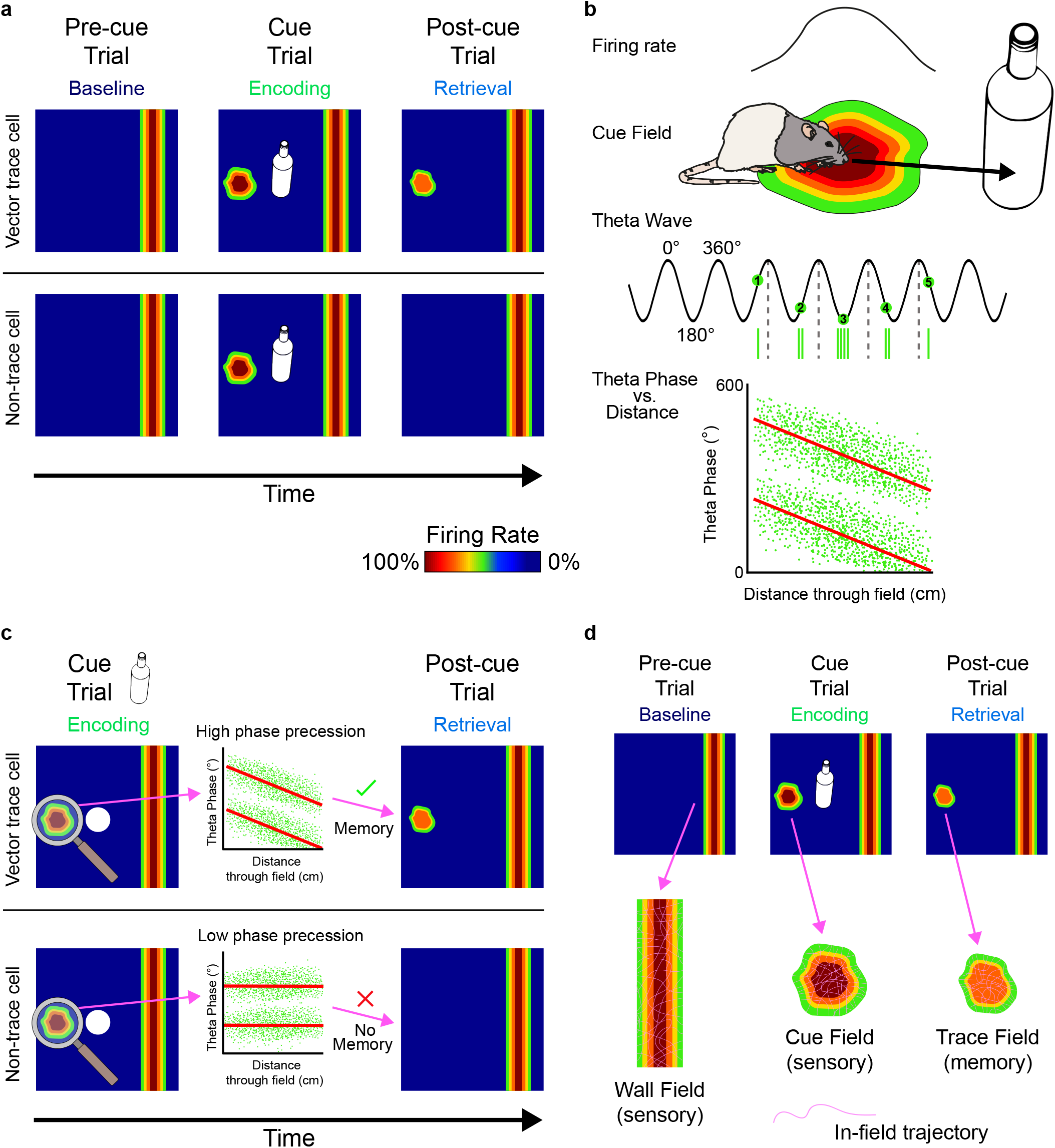
Testing ideas regarding theta phase precession for vector trace memory. a) Rats foraged for food in a 1m2 box while neurons were recorded in the dorsal subiculum. Heat-maps show firing rates of a neuron as a function of the position of the rat (warmer colours indicate higher firing rates). The three-trial series consisted of: 1) the Pre-cue trial; 2) the Cue trial, during which an object such as a barrier or wine bottle was inserted into the box; 3) the Post-cue trial, during which the cue was removed. We divided responses to cue removal into two types, according to the presence (top row: ‘Vector trace cell’) or absence (bottom row: ‘Non-trace cell’) of vector trace memory as determined from the Post-cue trial (Poulter et al, 2021). In both cases, inserting a cue elicited a new firing field (‘cue field’) additional to that of the wall field; however, a cue field could be subsequently forgotten (a ‘non-trace cue field’) or subsequently remembered (a ‘later-trace cue field’). **b**) Theta phase precession of in vector cells. A vector cell fires as the rat encounters an object at a preferred distance, e.g. 30cm, and allocentric direction, e.g. east, forming the cell’s ‘Cue field’. The cell fires at successively earlier phases of theta oscillation (sinusoidal trace), starting at a late phase (e.g. ∼280° for spike in first theta cycle) when the rat enters, and ending at an earlier phase (e.g. ∼70° in last theta cycle) as the rat leaves, the firing field. Vertical green bars = Spikes. Numbered green circles = average burst times in successive theta cycles. A phase-vs-distance scatterplot (bottom) can then be constructed from multiple traversals through the field. **c**) We addressed question 1, ‘Does phase precession enhance memorability?’ by asking if the proportion of phase precession was higher in ‘later-trace cue fields’ which were subsequently remembered than ‘non-trace cue fields’ which were subsequently forgotten. **d**) We addressed question 2, ‘Does phase precession occur in memory-dependent as well as sensory-driven fields?’ by comparing the proportions of phase precession in trace fields to those in wall fields and cue fields. A phase-vs-distance scatterplot for each field was constructed from all acceptable trajectories (pink lines) through the field. See Methods for the criteria defining acceptable trajectories.

### Theta phase precession: largely sensory-driven and/or absent in memory-dependent firing fields?

Intense research effort has gone into the understanding the mechanisms of theta phase precession (e.g. Mehta et al, 2002; Harris et al, 2002; Huxter et al, 2003; Zugaro et al, 2005; Harvey et al, 2009; Ravassard et al 2013; Aghajan et al, 2015), which remain much debated (reviewed e.g.: Buzsaki, 2006; Maurer and McNaughton, 2007; Burgess and O’Keefe, 2011; Colgin, 2013; Jaramillo and Kempter, 2017). Despite this attention, one question that has received little empirical study is to what extent phase precession is dependent upon activity that can be broadly characterised as *sensory-driven* vs *memory-driven*. Again, this is perhaps because it has been difficult to find unambiguous mnemonic correlates of hippocampal cell firing, and, correspondingly, to distinguish memory-dependent from sensory-driven activity. In our view, Subicular vector cells provide a useful model here.

The insertion of a cue object lawfully elicits a new firing field at an expected distance and direction from the cue object predicted by the Boundary Vector cell model (Hartley et al, 2000; Lever et al, 2002; Barry et al, 2006; Lever et al, 2009). We can thus characterise this new firing field as ‘sensory driven’. In some cases, when the cue object is now removed, that firing field persists (typically at lower rates and with some locational drift) in the form of a trace field (Poulter et al, 2021). We thus characterise this trace field as ‘memory-dependent’. We have speculated that cue fields, but not wall fields, are likely cell-specific zones for plasticity (Poulter et al, 2021). If so, perhaps the most useful baseline comparator for ‘memory-dependent’ trace fields are wall fields, since in our recording paradigm, walls are constantly perceptually present, while cue objects are not. Below we compare phase precession in trace fields (memory-dependent fields, Fig1d, right) to both cue fields and wall fields (types of sensory-driven fields Fig1d, left, middle).

Up to now, one line of evidence speaking to the sensory-vs-mnemonic activity question has come from depriving CA1 of either of its two key input regions, Entorhinal cortex and CA3. Interestingly, sharply-contrasting consequences for CA1 emerge from such deprivation. While lesioning medial entorhinal cortex (Schlesiger et al, 2015) profoundly disrupts CA1 theta phase precession, silencing CA3 (Davoudi and Foster, 2019) has no effect upon CA1 phase precession. (Instead, silencing CA3 profoundly disrupts sharp wave ripple activity (Davoudi and Foster, 2019), and thus presumably ripple-based replay.) Furthermore, in a correlational study that explicitly compared Entorhinal vs CA3 inputs to CA1 theta phase precession, stronger phase precession was associated more with Entorhinal-driven than CA3-driven activity (Fernandez-Ruiz et al, 2017). Taken together then, broadly characterising entorhinal cortex as conveying sensory input, and CA3 as conveying mnemonic input, it could be thought that theta phase precession is largely sensory-driven and thus potentially absent in memory-dependent firing fields. Accordingly, we were motivated to address this issue by asking our second question; *does theta phase precession occur* not only at the encoding stage (cue-elicited firing fields), but also *at the retrieval stage of memory, in the trace fields of vector trace cells*? If it does, then the much-theorised computational advantages of phase precession likely apply to memory-dependent fields, and thus likely support hippocampal memory-dependent functions such as episodic memory, imagination, and prospection (Byrne et al, 2007; Hassabis et al, 2007; Schacter et al, 2007).

## Results

### The presence of theta phase precession predicts memorability

#### Q1: ‘Do cue-elicited fields which are subsequently remembered show higher rates of theta phase precession than cue-elicited fields which are subsequently forgotten?’

In the (Poulter et al, 2021) paradigm, the four box walls were always present, and the vast majority of trials including training and pre-baseline trials contained no cue objects. Thus, cue objects, which varied, may have elicited a mismatch novelty signal directing new learning. We tested the hypothesis that theta phase precession would enhance such learning. Consistent with this idea, we found that subsequently-remembered cue fields (‘later-trace cue fields’) exhibited higher rates of phase precession (71%) than subsequently-forgotten cue fields (‘no-trace cue fields’) i.e. those which did *not* result in a trace field (49%) [*Distance:* Later trace: 48/68; No trace: 100/205, n = 273, χ^2^(1) = 9.78, p = 0.0018 (*Normalised Distance:* Later trace: 48/68; No trace: 102/205, χ^2^(1) = 8.95, p = 0.0028)].

In other words, the presence of phase precession positively predicted the memorability of the cue field. We call this the ‘phase precession memorability effect’.

Figure 2 shows three examples of the *presence* of phase precession in subsequently-*remembered* cue fields. Figure 3 shows three examples of the *absence* of phase precession in subsequently-*forgotten* cue fields.

**Fig. 2.**
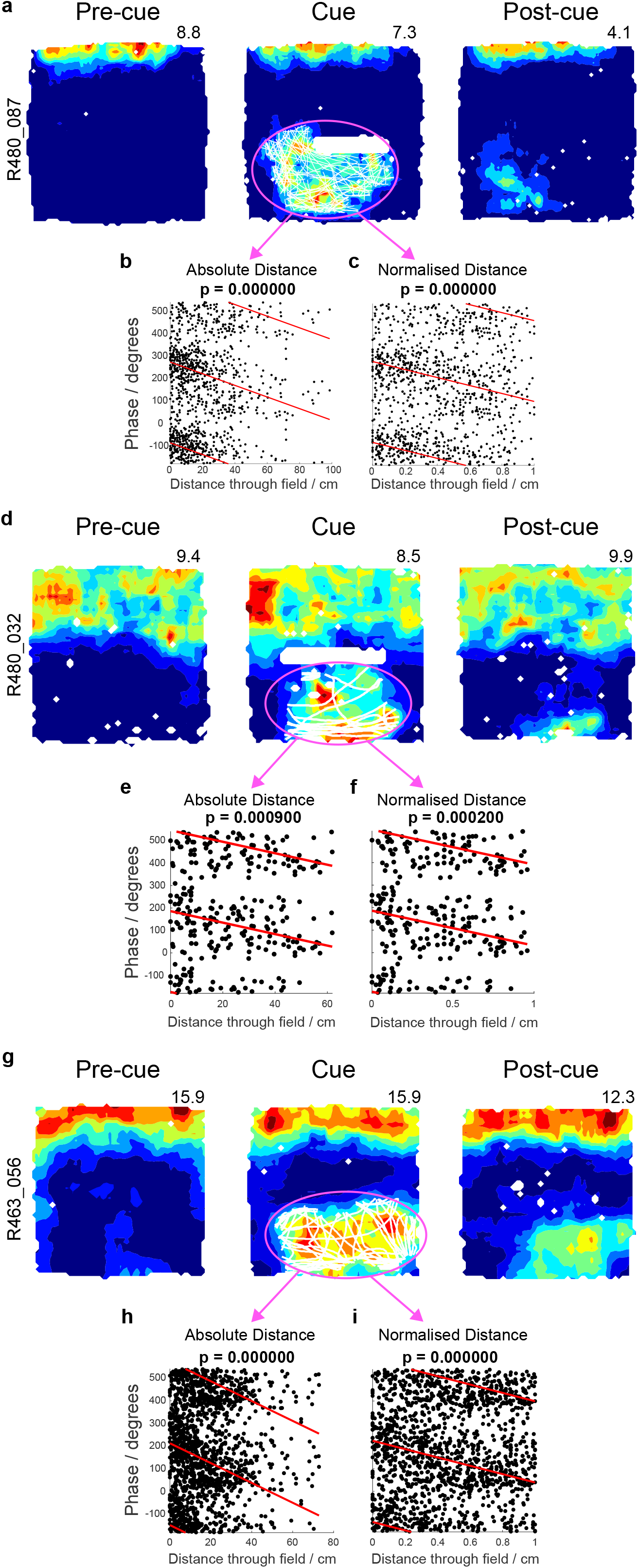
Three examples where theta phase precession is present in cue fields which are subsequently remembered (‘later-trace cue fields’) **a**) Example of a vector trace cell showing strong theta phase precession (as assessed by both absolute (**b**) and normalised (**c**) distance) in its cue field (circled pink), elicited by insertion of an object into a previously empty environment. Following object removal (Post-cue), the memory of the cue field persists in the form of a trace field. A second (**d**,**e**,**f**) and third (**g**,**h**,**i**) vector trace cell showing the same phenomenon. Rat and cell identifier numbers are shown on the left, and peak firing rate (Hz) at the top-right, of each firing-rate heat map.

**Fig. 3.**
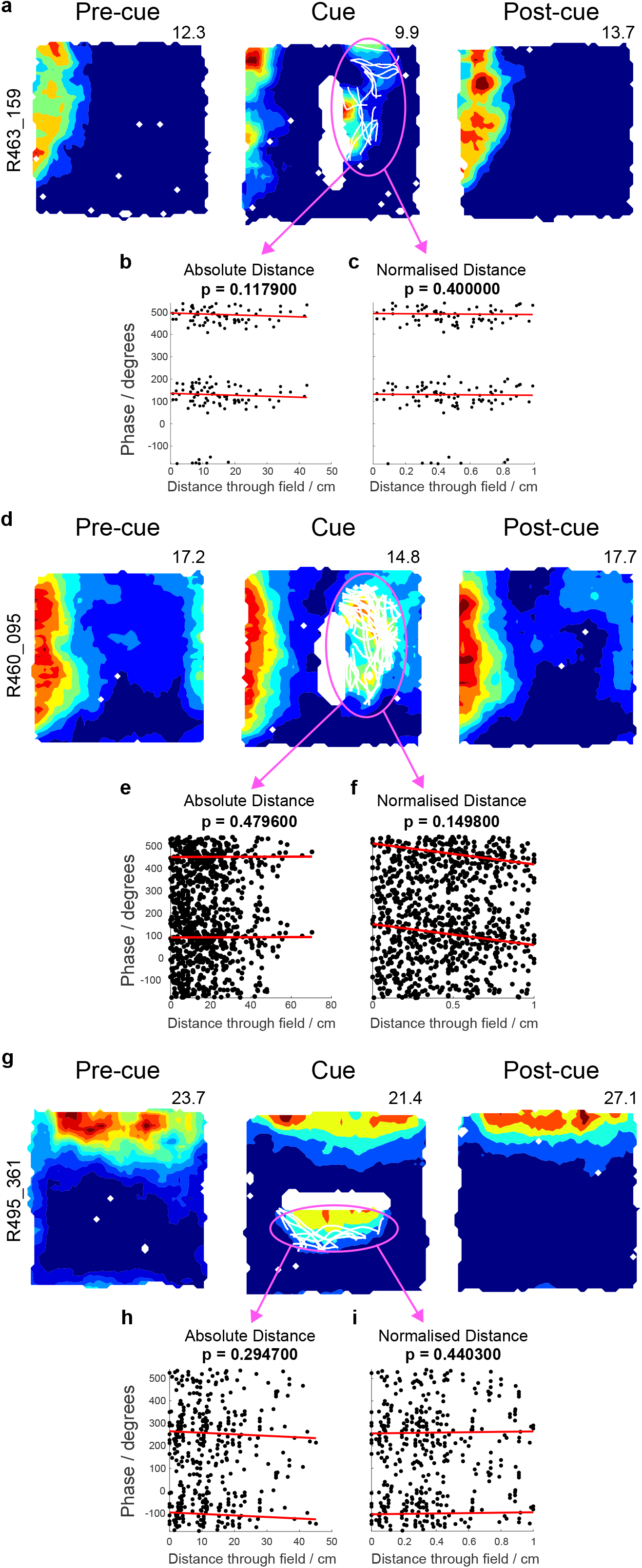
Three examples where theta phase precession is absent in cue fields which are subsequently forgotten (‘no-trace cue fields’) **a**) Example of a non-trace vector cell showing absent theta phase precession (as assessed by both absolute (**b**) and normalised (**c**) distance) in its cue field (circled pink), elicited by insertion of an object into a previously empty environment. Following object removal (Post-cue), the cue field is subsequently forgotten. A second (**d**,**e**,**f**) and third (**g**,**h**,**i**) non-trace vector cell showing the same phenomenon. For these two further examples, phase distribution is more dispersed than that shown in the first cell. Rat and cell identifier numbers are shown on the left, and peak firing rate (Hz) at the top-right, of each firing-rate heat map.

### The ‘phase precession memorability effect’ is not merely attributable to anatomical location within the subiculum

Poulter et al (2021) found that vector trace cells are more common, and theta modulation is somewhat higher, in the distal than proximal subiculum. Moreover, our analysis here shows that theta phase precession was more common in distal-recorded cue fields (59%) than proximal-recorded cue fields (38%) (*Distance:* Distal: 128/218; Proximal: 20/52, n = 270, χ^2^(1) = 6.95, p = 0.008). Interestingly, this *distal > proximal* relationship in subicular theta properties mirrors that in entorhinal cortex (*medial > lateral*, e.g. Deshmukh et al, 2010), consistent with reciprocal connectivity between distal subiculum and *medial* entorhinal cortex, and between proximal subiculum and *lateral* entorhinal cortex (anatomical evidence summarised in Poulter et al, 2021).

Accordingly, we re-ran analysis, on distal-recorded cue fields only, to ask if the ‘phase precession memorability effect’ extended beyond distal-vs-proximal biases. The results were similar: within the distal subiculum itself, subsequently-remembered cue fields exhibited higher levels of phase precession (73%), than subsequently-forgotten cue fields (56%), (*Distance:* Later trace: 46/63; Non trace: 87/156, n = 219, χ^2^(1) = 5.60, p = 0.018). Thus, the ‘phase precession memorability effect’ is not merely attributable to anatomical location within the subiculum.

### Theta phase precession in memory-driven trace fields

#### Q2: ‘Do memory-dependent vector trace fields show rates of theta phase precession which are comparable to sensory-driven vector fields?

To address this question, we compare trace fields (i.e. memory-dependent fields) with fields elicited by both inserted cues and by box walls. To increase statistical power, we included all trace fields in the (Poulter et al, 2021) dataset, i.e. including those not only from the post-cue trial that directly followed the cue trial, but also from repeated post-cue trials (where these were run). Figure 4 presents examples from two vector trace cells, showing repeated observations of phase precession in trace fields. Post-cue trials 1,3,5 are shown for one cell (Figure 4A-C), and post-cue trials 1 & 2 for the other (Figure 4D-F), where the last post-cue trials shown were the last post-cue trials tested for each cell respectively. Thus, it is possible that phase precession is quite long-lasting in trace fields, though this clearly warrants further study.

**Fig. 4.**
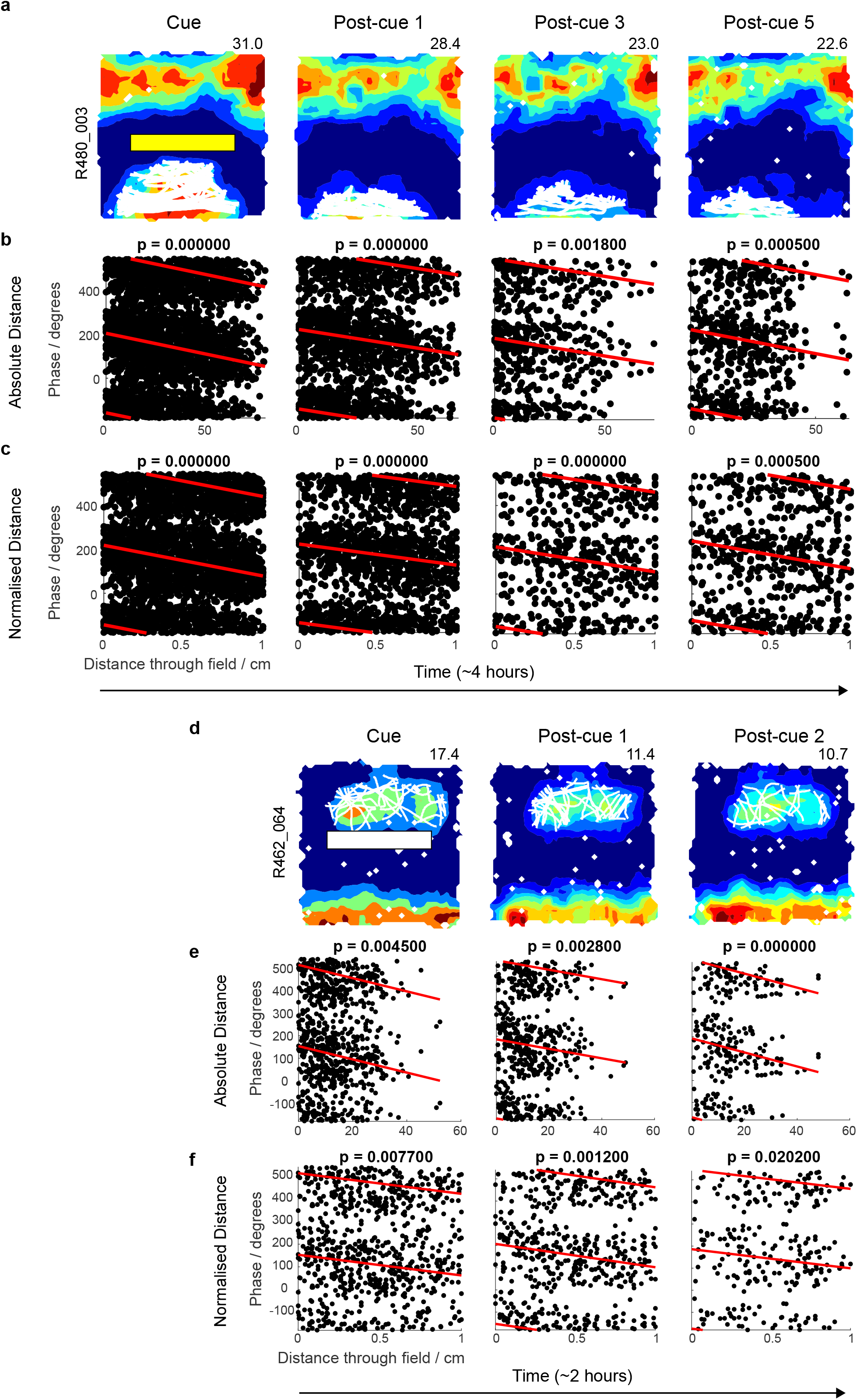
Theta phase precession is robust in trace fields. Two examples of vector trace cells [cell 1 (**a**-**c**); cell 2 (**d**-**f**)], where theta phase precession occurs in cue fields and trace fields, and persists as long as tested in trace fields. This suggests that the theorised computational benefits of theta phase precession for cognition extend to memory-dependent vector representations. Post-cue trial 5 for cell 1, and post-cue trial 2 for cell 2, were the last post-cue trials run for each cell. Rat and cell identifier numbers are shown on the left, and peak firing rate (Hz) at the top-right of each firing-rate heat map.

#### Comparing trace fields with sensory-driven cue fields

Despite having relatively small spatial extent, over a third of trace fields showed theta phase precession [*Distance:* 36%: 33/92; (*Normalised Distance:* 39%: 36/92)]. This proportion was lower than that in cue fields, where over half showed theta phase precession [*Distance:* Cue fields 54% (148/273), vs trace fields: χ^2^(1) = 9.26, p = 0.002; (*Normalised Distance:* 55% (150/273, vs trace fields: χ^2^(1) = 6.89, p = 0.009)]. The higher proportion of theta phase precession in cue fields than trace fields is consistent with the possibility that, at least in the (Poulter et al, 2021) paradigm, the cue field was a privileged zone for enhanced encoding, and that, as suggested above, phase precession might function to enhance learning.

#### Comparing trace fields with sensory-driven wall fields

We next compared trace fields to a different set of sensory-driven fields, namely wall fields. In our paradigm, the four box walls remained throughout the entire experiment and thus were constantly available to perception (e.g. visual, somatosensory). To increase statistical power, we included all wall fields in the (Poulter et al, 2021) dataset.

Remarkably, the proportion of trace fields with phase precession was very similar to that of wall fields (*Distance:* Trace fields: 36%, Wall fields: 35%; χ^2^(1) = 0.01, p = 0.91; *Normalised Distance:* Trace fields: 39%; Wall fields: 36%, χ^2^(1) = 0.31, p = 0.57; trace field n = 92; wall field n = 1538; Total n = 1630). Accordingly, we conclude that theta phase precession is at least as robust in memory-driven trace fields as it is in sensory-driven wall fields.

## Discussion

Theta phase precession is a complex, emergent phenomenon; it is not trivial to selectively suppress it without affecting constituent factors. Here, we explored already-present variance in vector trace memory in the (Poulter et al, 2021) dataset, and found that theta phase precession in a cue field in the encoding trial (cue trial) was associated with an increased likelihood that the cue field would be subsequently remembered as a trace field in the retrieval trial (post-cue trial), following removal of that cue. This increased likelihood was still observed when analysis was restricted to the distal subiculum, where theta modulation is generally higher than in proximal subiculum. This result is consistent with the possibility that phase precession is not simply a physiological marker of a cell’s anatomical location, but may actually enhance vector field encoding.

Exactly how theta phase precession might enhance encoding in vector trace cells remains to be elucidated. In the Introduction, we presented the consensus view that phase precession may enable learning of spatial and temporal sequences. Briefly, we consider this further here. In some theta cycles in the cue trial (encoding trial), the spikes representing the cue fields of subicular vector trace cells will occur towards the middle/end of these sequences. It seems reasonable to hypothesise that trace fields may be encoded by repetitions of such sequences, whereby cells (of various types) firing earlier on in the sequence, under conditions favouring long-term plasticity, come to activate the cells representing cue fields in a predictive manner, and thus even when the cues that elicited those cue fields are no longer present. The sets of cells which, subsequent to plasticity, trigger vector trace cells to fire could represent, say, paths towards the cue fields, movement direction, and global spatial context. Such a process is likely to involve subsequent ‘replay’ of the sequences embedded in repeated theta cycles (Drieu et al, 2018), especially if, as we now suspect, trace fields can last for many hours and days. We have little data to constrain hypotheses about the potential CA3/CA1 and subicular networks involved in generating trace fields in vector trace cells. However, we note that: 1) the subiculum can generate ripples independently of CA3-CA1 propagation (Imbrosci et al, 2021); 2) relatedly, subicular pyramidal cells show high pyramidal-pyramidal cell recurrent connectivity (perhaps even higher than in CA3) (Bohm et al, 2018). Thus, we cannot exclude the possibility that purely intra-subicular networks could generate trace fields via theta-to-ripple sequence consolidation. In this respect, we note that the role of theta cycle length has not been much considered. We recently proposed a Novelty-Elicited Slowing of Theta hypothesis (Hines et al, 2022), positing that hippocampal theta frequency slows in novelty, so theta cycles lengthen, in order to increase items-per-cycle and memorability, as part of a general ‘need-for-encoding’ process. We suggest that ‘need-for-encoding’ apparently favours the single theta cycle as a privileged encoding packet in the trade-off between more-memoranda-per-packet (A->B->C->D, *longer* cycles preferred) vs more repetitions (A->B->C, B->C->D, *shorter* cycles preferred).

Irrespective of the contribution of theta phase precession to the memorability of cue fields, an additional observation was that theta phase precession could be seen in over a third of the trace fields of vector trace cells. We speculate that this would be a high enough proportion for subicular networks relying upon phase precession in vector trace cells to support hippocampal memory-dependent functions such as episodic memory, imagination, and prospection (Byrne et al, 2007; Hassabis et al, 2007; Schacter et al, 2007). Interestingly, high-resolution fMRI mapping has implicated the subiculum as part of the default mode network (DMN), in particular part of DMNa, which is coupled to the retrosplenial cortex and parahippocampal cortex (Braga et al, 2019). Thus, it may be that, for instance, spatial relationships between elements in a scene could be decoded from theta phase relationships between vector trace cells, and particularly amenable to DMN-type cognitive operations.

## Methods and Materials

This study further analysed data in the (Poulter et al. 2021) dataset. Accordingly, some of the methods and materials in that report are reproduced here.

### Subjects

Six male Lister hooded rats, weighing 392–522 g (aged 3–5 months) at the time of surgery, were used as subjects. All rats were maintained on a 12–12-h light–dark schedule (with lights off at 10:00; all animals were tested during their dark phase). Food deprivation (after rats had recovered from surgery) was maintained during recording periods such that subjects weighed 85–90% of free feeding weight. All experiments were performed under the Animals (Scientific Procedures) Act 1986. Approval for the animal experiments was granted by both Durham University AWERB and the UK Home Office Project and Personal Licenses.

### Surgery and tetrode implants

Under deep anesthesia (1–3% isoflurane) and using intra- and post-operative analgesia (buprenorphine, 0.04 mg per kg), rats were chronically implanted with two microdrives (one above the dorsal subiculum of each hemisphere). These microdrives allowed four or eight tetrodes to be vertically lowered through the brain. The eight-tetrode-loaded microdrives (implanted in four rats) used custom three-dimensionally-printed barrels to create a 4 × 2 tetrode array (150-μm spacing between barrel holes). With the four-tetrode-loaded microdrives (implanted in two rats), all the tetrodes were loaded using a single cannula. Tetrodes were constructed from HM-L-coated platinum–iridium wire (90%/10%, California Fine Wire, 25 μm). The details of tetrode mapping for each drive were recorded using photographs and notes both before surgery and after perfusion.

### Subiculum: implant co-ordinates and histology

Our implants targeted the anterior portion of the dorsal subiculum. The skull coordinates used for insertion of the centroid of the tetrode array were based on Paxinos & Watson 2007 in the following range: anterior–posterior (AP):−5.8 to −6.4 mm; medial–lateral (ML): ± 2.9–3.3 mm. Details of recording sites were reconstructed using records of electrode movement, physiological markers and post-mortem histology. The rats were killed and transcardially perfused with saline followed by 4% paraformaldehyde. Each brain was coronally sliced into 40-μm thick sections, mounted and Nissl-stained (using cresyl violet or thionin) for visualization of the electrode tracks/tips. Digital photomicrographs were converted from color to black and white, and contrast and brightness adjusted, using Photoshop Express 3.5. Data from recording sites in the CA1 or the dorsal presubiculum were excluded, and recording sites in the subiculum were classified as being located in either the proximal subiculum or the distal subiculum (see ref. Poulter et al. 2021 for detailed methodology of how the dorsal subiculum was parcellated into proximal and distal zones).

### Electrophysiological recording

Rats were allowed 1 week to recover post-operatively before screening sessions began. During screening and inter-trial intervals, the rat rested on a square holding platform (40-cm sides, 5-cm high ridges) containing sawdust. Tetrodes were gradually lowered toward the subiculum pyramidal layer over days/weeks. Tetrodes were left to stabilize for at least 24h after tetrode movement before recording commenced. Electrophysiological data from screening and recording sessions were obtained using Axona DACQ systems (DacqUSB). Electrode wires were AC-coupled to unity-gain buffer amplifiers (headstage). Lightweight wires (∼4 m) were connected the headstage to a pre-amplifier (gain 1,000). The outputs of the pre-amplifier passed through a switching matrix, and then to the filters and amplifiers of the recording system (Axona). Signals were amplified (6,500–14,000) and band-pass filtered (500 Hz to 7 kHz). Each channel was continuously monitored at a sampling rate of 50 kHz, and action potentials were stored as 50 points per channel (1 ms, with 200-μs pre-threshold and 800-μs post-threshold) whenever the signal from any of the 4 channels of a tetrode exceeded a given threshold. LFP signals were amplified 3,500–5,000, band-pass filtered at 0.34–125 Hz and sampled at 250 Hz. Two arrays of infrared light-emitting diodes (LEDs), one array larger than the other for tracking discriminability, were attached to the head of the rat to track head position and orientation using a video camera and tracking hardware/software (DacqUSB, Axona). The arrays of LEDs were positioned such that the halfway position between the two arrays was centered above the skull of the rate. Offline analysis defined this halfway position as the position of the rat (TINT, Axona). Positions were sampled at 50 Hz.

### Testing laboratory and recording environments

External cues such as a lamp, PC monitor and cue cards on the walls provided directional constancy throughout the test trial series. For every trial, the rat was carried directly from the holding platform with its head facing toward the recording arena. During trials, the rat searched for grains of sweetened rice randomly thrown into the box about every 30 s. At the end of each trial, the rat was removed from the recording environment and placed back on the holding platform until the next trial. Inter-trial intervals varied from 10 min to 1 h, but were typically around 25 min. The standard recording environment was a square box (100 × 100 × 50-cm high) painted in ‘light rain’ gray. Occasionally, for more distally tuned cells, a larger, same colored environment was used (either 150 × 150 × 50-cm high or 150 × 190 × 50-cm high). Four types of cue, introduced into the box during the cue trial, were used: a black painted barrier (50 × 2.5 × 50-cm high); three wooden black bricks juxtaposed along their long axis (20 × 9.5 × 4.5-cm high), thus creating a 60 × 28.5 × 4.5-cm high cue; a high-contrast white stencil-painted stripe (60 × 10 × 0-cm high) acting as a purely visual cue; and wine bottles (base diameter 7 cm, 30-cm high) painted with different large, high-contrast patterns and/or affixed with somatosensorily different patches. In some trials, only one bottle was inserted into the environment. In other trials, two or more bottles were inserted in different configurations: tightly juxtaposed in an array to create a continuous barrier, placed apart to create a linear spaced array or placed apart in different regions of the box.

### Standard test trial sequence and variants

The standard test trial sequence consisted of three consecutive trials: (1) a pre-cue trial, in which the recording box contained no cue; (2) a cue trial, in which the box contained one of the abovementioned four types of cue; and (3) a post-cue trial, in which the box again contained no cue. In a minority of sessions, one variant of the standard test sequence was that more than one cue trial was run successively before the post-cue trial. In this case, only the first cue trial was used to define the cue field and cue responsiveness of the neuron. This procedure had no significant effect on the probability of a trace response arising (successive multiple cue trials 42% trace, single cue trials of matched cue type (barrier), 33% trace; Z-test for proportions: Z = 0.84, P = 0.40). In other cases, the experimental session was extended to include a repeat of the standard three-trial sequence (pre-cue trial, cue trial, post-cue trial), most often with physically different cues. In these cases, only the single cue trial with the strongest cue-elicited field (that is, highest integrated firing rate, see description below) and its accompanying pre-cue and post-cue trials were selected for main analysis.

### Cell isolation

Cluster cutting to isolate single units was performed using a combination of KlustaKwik (v.3)62 and manual isolation using TINT (Axona). All the trials from a given session were loaded into TINT as a merged dataset, which was clustered using KlustaKwik’s principal component analysis. Subsequent manual adjustments were made where necessary. Once merged-trial cutting was complete, cell clusters were automatically split into each individual trial of that session (Axona MultiCutSplitter).

### Firing-rate maps

Firing-rate maps for all recorded neurons were produced by first dividing the recording arena into a grid of 2 × 2-cm square spatial bins and finding the summed occupancy time and number of spikes fired in each bin. Summed occupancy and spiking maps were then smoothed with a 10 × 10-cm boxcar kernel, and rate maps were constructed by dividing summed spiking by summed occupancy. Data from periods of immobility (movement speed <5 cm s−1) were excluded from rate maps, so as to restrict the analysis to neural firing from theta epochs, and exclude firing occurring during sharp-wave ripple epochs.

### Definition of cue fields and cue-responsive neurons

New firing fields generated by the insertion of a cue were detected as follows. First, firing-rate maps were converted to z-scores to allow comparison between different trials, even following trial-to-trial fluctuations in the firing rate. For each map, values for each bin had the overall mean firing rate subtracted and were then divided by the overall variance of the firing rate across bins. Following this, the z-scored pre-cue map was subtracted from the z-scored cue map, thus producing a map describing where firing was increased specifically during the cue trial relative to the pre-cue trial. Cue fields were then defined as contiguous regions of the resulting map with a value of ≥1. If more than one cue field was present, only the largest was used for further analysis. Cue-responsive cells were defined as those where the sum of z-scored firing rate, within the cue field, was ≥70.

### Definition of wall fields

Baseline firing fields in response to the walls of the environment were detected as follows. First, firing-rate maps were converted to z-scores to allow comparison between different trials, even following trial-to-trial fluctuations in firing rate. For each map, values for each bin had the overall mean firing rate subtracted and were then divided by the overall variance of the firing rate across bins. Following this, the z-scored rate map from the most recent cue-less trial (e.g. the last pre- or post-cue trial) was subtracted from the trial in question with this difference thresholded at zero to contain only non-negative values. Finally, these non-negative values were subtracted from the z-scored rate map for the trial in question. Wall fields were then defined as contiguous regions of the resulting map with a value of ≥1, where the sum of z-scored firing rate within the wall field was ≥70. If more than one wall field was present, only the largest was used for further analysis.

### Definition of trace and overlap scores

To define trace and overlap scores, cue and post-cue firing-rate maps were z-scored, and the pre-cue map was subtracted from both, so as to highlight changes in firing relative to the pre-cue trial (the same procedure as described above, “Definition of cue fields and cue-responsive neurons”). The trace score was defined as the mean value of z-scored firing within the cue field region (as defined above) in the post-cue trial, divided by the mean of z-scored firing within the cue field region in the cue trial. A trace score of 1 therefore indicates a memory-based response of equal strength to that induced by the presence of the cue. To define the overlap score, we first detected whether any new regions of firing existed in the post-cue trial, relative to the pre-cue trial: such post-cue fields were defined as contiguous regions of the (z-scored, pre-cue subtracted) post-cue map with a value of ≥1. If several post-cue fields existed, only the largest was used for further analysis. If a post-cue field was present, the overlap score was defined as follows:

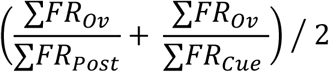

Where ∑*FR* in all cases refers to the summed firing rate in a region of the post-cue trial rate map: ∑*FR*_*post*_ being the summed rate in the post-cue field, ∑*FR*_*Cue*_ the summed rate in the cue field, and ∑*FR*_*Ov*_ being the summed rate in the overlap between the cue and post-cue fields. Overlap score therefore assesses the average extent to which post-cue field firing overlaps the cue field firing, and vice versa. Where no post-cue field was present, the Overlap score was set to zero.

### Theta-firing-phase analysis

Instantaneous theta phase was defined by filtering LFPs using a 5–11-Hz band-pass filter and taking the angle of the Hilbert-transformed filtered signal. The theta phase of each spike was defined as the phase of the temporally corresponding LFP sample, from the hemisphere-matched LFP in the distal subiculum with the highest signal-to-noise ratio for the theta oscillation. The signal-to-noise ratio for theta was defined as the mean power in the theta band (±0.5 Hz around the highest power between 7 and 10 Hz) divided by the mean power in the range 2–20 Hz, excluding the theta band. Spectral power was estimated using the fast-Fourier transform. The overall theta modulation of each neuron was estimated using the length of the resultant mean vector of phases. As with rate maps, speed filtering (>5 cm s−1 epochs passing) was applied; only neurons firing ≥50 spikes while the animal occupied the relevant firing field (over the course of a whole trial), and with significant theta phase modulation (defined as Rayleigh test P < 0.01), were used for phase analysis. This was done to remove the influence of noisy data. The preferred firing phase of each neuron, in each firing field, was defined as the circular mean of the spike phases for spikes occurring while the rat was within the given firing field (wall field, cue-field regions). The above steps were performed using custom-written Matlab scripts.

### Theta phase precession analysis including pre-analysis

Runs through the wall, cue and trace fields, defined above, were detected and trajectories meeting the following criteria being used for subsequent phase precession analyses: path length > 10cm; minimum speed > 5 cm s−1; mean speed > 10 cm s−1; and tortuosity < 1.5 (tortuosity defined as path length divided by the Euclidean distance between field entry and exit points). Absolute distance through field was measured as the path length along the trajectory, with this being divided by the total path length of the trajectory in the Normalised case. Only neurons firing ≥ 10 spikes across these filtered trajectories were used in the subsequent phase precession analyses. Circular-linear correlation coefficients between the theta phase of spikes and the distance travelled through a field were calculated using equation 3 in Kempter et al., 2012. Fields were considered to show significant phase precession if the correlation coefficient was more negative than 95th percentile of the shuffled distribution of phase versus distance travelled (N=10000 shuffles)

